# Laminin-511 and α6 Integrins Regulate the Expression of CXCR4 to Promote Endothelial Morphogenesis

**DOI:** 10.1101/846022

**Authors:** Hao Xu, Kevin Pumiglia, Susan E. LaFlamme

**Affiliations:** Department of Regenerative and Cancer Cell Biology, Albany Medical College, Albany NY 12208

**Keywords:** Integrin, laminin, CXCR4, angiogenesis

## Abstract

During angiogenesis, endothelial cells engage components of the extracellular matrix through integrin-mediated adhesion. Endothelial expression of laminin-411 and laminin-511 are known to promote vessel stability. However, little is known about the contribution of these laminins to endothelial morphogenesis. We used two organotypic cell culture angiogenesis assays in conjunction with RNAi approaches to demonstrate that depletion of either the α4 chain of laminin-411 or the α5 chain of laminin-511 from endothelial cells inhibits sprouting and tube formation. Depletion of α6 integrins resulted in similar phenotypes. Gene expression analysis indicated that loss of either laminin-511 or α6 integrins inhibited the expression of CXCR4, a gene previously associated with angiogenic endothelial cells. Pharmacological or RNAi-dependent inhibition of CXCR4 suppressed endothelial sprouting and morphogenesis. Importantly, expression of recombinant CXCR4 rescued endothelial morphogenesis when the α6 integrin expression was inhibited. Additionally, the depletion of α6 integrins from established tubes resulted in the loss of tube integrity and laminin-511. Taken together, our results indicate that α6 integrins and laminin-511 can promote endothelial morphogenesis by regulating the expression of CXCR4 and suggest that the α6-dependent deposition of laminin-511 protects the integrity of established endothelial tubes.

**Summary statement:** Endothelial-secreted laminin-511 and α6 integrins promote endothelial morphogenesis by regulating the expression of the chemokine receptor, CXCR4. The depletion of α6 integrins from established tubes results in the loss of tube integrity and laminin-511.

## Introduction

Angiogenesis is a process by which new vessels sprout from the pre-existing vasculature, anastomose with neighboring sprouts to form new networks, which then remodel into mature vessels (Carmeliet, 2003). Angiogenesis contributes to both normal and pathological processes. Angiogenesis associated with tissue repair, tumor progression, and inflammation share many of the same molecular mechanisms (Carmeliet, 2005). Therefore, gaining a better understanding the mechanisms that regulate various aspects of this process remains an important objective.

During early stages of angiogenesis, endothelial cells interact with components of the extracellular matrix (ECM), such as those present in the provisional matrix that is formed during tissue repair (Eming et al., 2007). Although endothelial cells can adhere to matrix proteins provided by other cell types, endothelial cells themselves secrete both ECM and basement membrane components, including fibronectin and the specific laminin isoforms that contribute to angiogenesis and stability respectively (Hallmann et al., 2005; Turner et al., 2017). Endothelial cells express two laminin isoforms, laminin-411 and laminin-511 (Hallmann et al., 2005). Mouse genetic studies have indicated that they both promote vessel stability (Song et al., 2017; Thyboll et al., 2002). However, the contribution of these laminins to endothelial morphogenesis has not been examined.

Integrins are α/β heterodimeric receptors that bind to ECM proteins, including components of the basement membrane to mediate adhesion and to activate signaling pathways that cooperate with growth factor receptors to regulate cell behavior (Danen and Sonnenberg, 2003; Streuli and Akhtar, 2009). Several endothelial integrin heterodimers including α5β1, αvβ3, α2β1, and α6β1 are known to regulate angiogenesis [reviewed in (Avraamides et al., 2008; Davis and Senger, 2005) and their individual roles can be context dependent (Avraamides et al., 2008; Davis and Senger, 2005; Turner et al., 2017). Alpha 6 integrins are receptors for laminin-411 and laminin-511 (Hallmann et al., 2005; Kortesmaa et al., 2000). Mouse genetic studies and *in vivo* antibody blocking experiments demonstrated that α6 integrins regulate angiogenesis by several mechanisms: the inhibition of the expression or function of α6 integrins on endothelial cells, endothelial progenitors, and macrophages (Bouvard et al., 2012; Bouvard et al., 2010; Bouvard et al., 2014; Germain et al., 2010; Primo et al., 2010 5; Seano et al., 2014).

Our current study employed organotypic cell-culture models to determine whether endothelial secreted laminin-411 and/or laminin-511 contribute to endothelial tubulogenesis in an ECM environment similar to the provisional matrix present in healing wounds (Eming et al., 2007;Bishop et al., 1999; Mavria et al., 2006; Nakatsu and Hughes, 2008; Nakatsu et al., 2003). We demonstrate that in these models endothelial cells form tubes that are associated with endothelial-secreted basement membrane components, including laminin-411 and laminin-511. RNAi-dependent inhibition of the expression of either the α4 chain of laminin-411 or the α5 chain of laminin-511 inhibited endothelial sprouting and tubulogenesis indicating that endothelial cells secrete these laminins to promote endothelial morphogenesis. Interestingly, inhibiting the expression of α6 integrins phenocopied the inhibition of laminin-411 and laminin-511. Previous *in vitro* cell culture studies employed either collagen gels or Matrigel to examine the role of α6 integrins in endothelial morphogenesis. These studies demonstrated that α6 integrins are required for endothelial cells to form cords in Matrigel, which is rich in laminin-111 (Primo et al., 2010). Interestingly, α6 integrins were not required in collagen gels (Primo et al., 2010). It is important to note that Matrigel does not support the formation of endothelial tubes (Simons et al., 2015). Thus, the organotypic cell culture models used in our current study provided the opportunity to understand the mechanisms by which α6 integrins and endothelial secreted laminins regulate tubulogenesis. We asked whether α6 integrins and their laminin ligands regulate the expression of genes known to regulate angiogenesis. We found that both α6 integrins and laminin-511 promoted the expression of the pro-angiogenic chemokine receptor CXCR4 (Salcedo and Oppenheim, 2003; Salvucci et al., 2002; Tachibana et al., 1998; Unoki et al., 2010). In addition, we show that pharmacological or RNAi-dependent inhibition of CXCR4 impaired sprouting and tubulogenesis and that the expression of recombinant CXCR4 in cells depleted of α6 integrins partially rescued the α6-knockdown phenotype. This is the first report to our knowledge that α6 integrins promote endothelial morphogenesis by regulating the expression of CXCR4. Lastly, we show that α6 integrins regulate the stability of endothelial tubes at least in part by regulating the endothelial deposition of laminin-511. Taken together, our data suggest that the interaction of α6 integrins with laminin-511 contributes to angiogenesis by regulating the expression of CXCR4 during endothelial morphogenesis, and that once tubes have formed α6 integrins are required for the deposition of laminin-511 and the stability of endothelial tubes.

## Results

### Endothelial laminins regulate endothelial tubulogenesis

To examine the contribution of endothelial laminin-411 and laminin-511 in tubulogenesis, we employed RNAi technology in conjunction with two organotypic angiogenesis assays: the planar co-culture model and the bead sprout assay. In the planar co-culture model, human endothelial cells (HUVECs) are plated at very low density on a confluent layer of human dermal fibroblasts (Bishop et al., 1999). Endothelial cells then proliferate and migrate to form cell trains/cords, which then form lumenized tubes over time (Bajaj et al., 2010; Bishop et al., 1999; Mavria et al., 2006). In the bead sprout assay, endothelial cells are adhered to cytodex beads and embedded in a fibrin gel, which is then covered with a confluent layer of human dermal fibroblasts. Endothelial cells sprout out from the beads into the fibril gel and form lumenized tubes over time (Nakatsu and Hughes, 2008; Nakatsu et al., 2003). Importantly, endothelial cells express laminin-411 and laminin-511 in addition to Col IV in both organotypic models as demonstrated by immunofluorescence microscopy (Fig. 1A-E).

**Figure 1.**
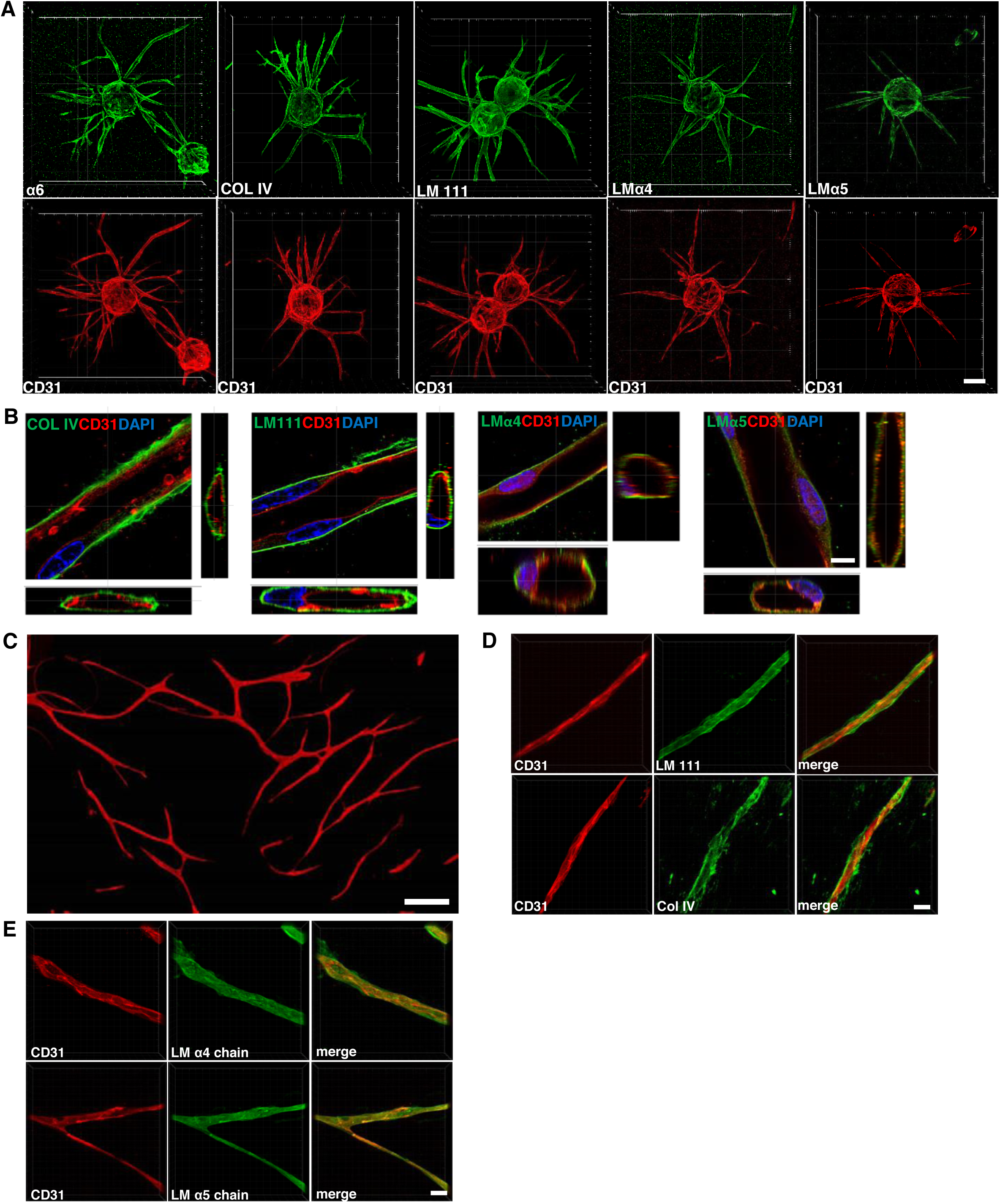
Laminin-411 and laminin-511 are deposited by endothelial cells in organotypic cultures. **(A)** Confocal images of 6-day bead sprouts immunostained for α6 integrins, Collagen IV, laminin (LM) β1 and γ1 chains, laminin α4 (LM-α4), and α5 (LM-α5) chains. Scale = 100 µm. **(B)** High magnification confocal images of lumenized sprouts and basement membrane components expressed on the basal surface. Scale = 6 µm. **(C)** Endothelial cells that have formed branch structures by day 6 in planar co-culture. Scale = 100 µm. **(D-E)** Confocal images of endothelial tubes formed in planar co-culture at day 3 immunostained for collagen IV, laminin (LM) β1 and γ1 chains, laminin α4 (LM-α4), and α5 (LM-α5) chains. Scale = 20 µm.

To determine whether the expression of either laminin-411 or laminin-511 was necessary for endothelial morphogenesis, we inhibited their expression with siRNA targeting either the α4 chain of laminin-411 or the α5 chain of laminin-511 and assayed the effects using both types of assays. Data was analyzed from three independent experiments. The efficiency of knockdown in each experiment was determined by qPCR (Fig. 2A-B). Depletion of either laminin chain inhibited sprouting in the bead-sprout assay (Fig. 2C-E). Both the number and lengths of sprouts were decreased (Fig. 2D-E). Similarly, knockdown of either LM-α5 or α4 chains caused defective endothelial morphogenesis in planar co-culture with a more dramatic phenotype resulting from inhibiting the expression of the α5 chain (Fig. 2F-G). Quantitation of the lengths of cell trains/cords formed after two days showed a mean length of 340 µm in control, whereas depletion of either the Lm-α4 or-α5 chain resulted in an average length of 175 µm and 95 µm, respectively. Our data indicate that both laminin-411 and laminin-511 contribute to endothelial sprouting and tube formation and that the expression of one isoform cannot compensate for the loss of the other suggesting that each laminin isoform may play distinct role during tubular morphogenesis.

**Figure 2.**
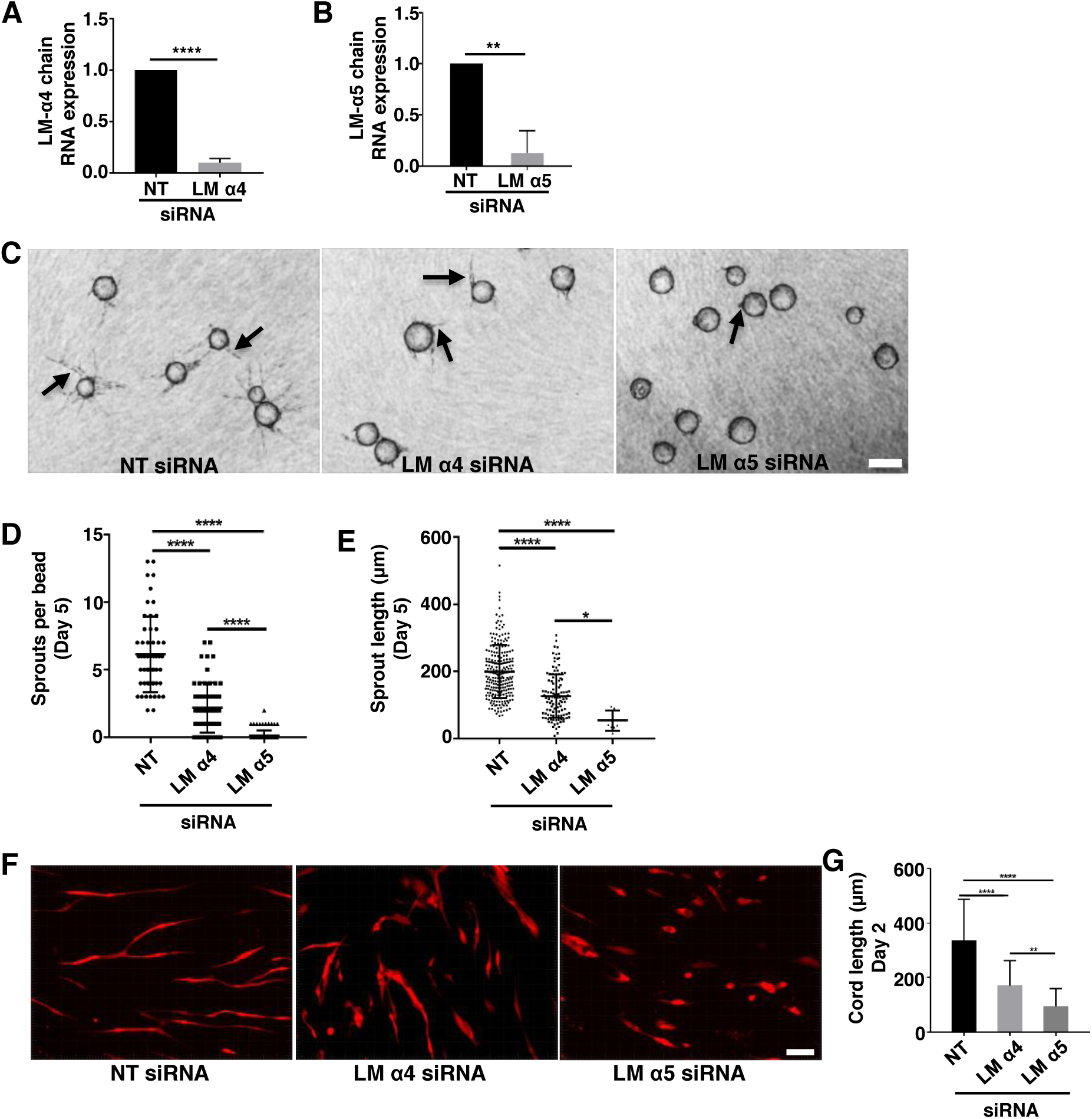
Laminin-411 and laminin-511 regulate endothelial tubular morphogenesis. **(A-C)** The expression of the LM-α4 or LM-α5 chain in three independent experiments was inhibited by siRNA. The efficiency of knockdown of the LM-α4 chain **(A)** and LM-α5 chain **(B)** in endothelial cells used in functional studies was assayed by qPCR. Plotted is the mean ± s.d. **(C)** Representative images of endothelial cells depleted of either the LM-α4 or -α5 chain in a 5-day bead sprout assay. Arrows indicate examples of sprouts. Scale = 250 µm. **(D-E)** Quantitation of number of sprouts **(D)** and sprout length **(E)** from 6-8 beads in each of 10 randomly selected fields in 3 independent experiments. Plotted are the **(D)** numbers of sprouts on individual beads or **(E)** or the length of individual sprouts. Data was analyzed by one-way ANOVA. The mean ± s.d. is indicated. **(F)** Representative images of RFP-expressing endothelial cells at day 2 of planar co-culture depleted of either the LM-α4 or LM-α5 chain compared to control. Scale = 100 µm. **(G)** Quantitation of endothelial cord length at day 2 for the different conditions. Plotted is length of endothelial cords measured from 10 random fields in 3 independent experiments. Data was analyzed by one-way ANOVA and plotted as the mean length ± s.d.

### Endothelial α6 integrins promote endothelial tubulogenesis

Alpha 6 integrins are known to bind to both laminin-411 and laminin-511 (Kortesmaa et al., 2000) and are expressed by endothelial cells in co-culture as expected (Fig. 3A). To determine whether depletion of α6 integrins phenocopied the loss of laminin-411 or laminin-511, we employed lentiviral vectors for the doxycycline inducible expression of either a non-targeting shRNA or one of three shRNAs targeting the integrin α6 subunit. It is important to note that the induction of these shRNAs results in the co-expression of a GFP reporter. Endothelial cells were transduced with these lentiviral vectors and the expression of shRNAs was induced by the addition of doxycycline. Effects of α6 depletion on endothelial morphogenesis in the planar co-culture were analyzed in three independent experiments. The efficiency of knockdown from these experiments was analyzed by western blot (Fig. 3B, left panel). Effects on endothelial cord/tube length were quantified every two days for a total of 10 days (Fig. 3B, middle panel). Images of representative co-cultures at day 6 are shown in Figure 3B (right panel). Taken together, the data show that morphogenesis is inhibited by day 2 and remained suppressed with very little lengthening of individual endothelial cords in the α6-depleted condition at day 10. Similar results were obtained with two additional shRNA targeting sequences (Supplemental Fig. 1).

**Figure 3.**
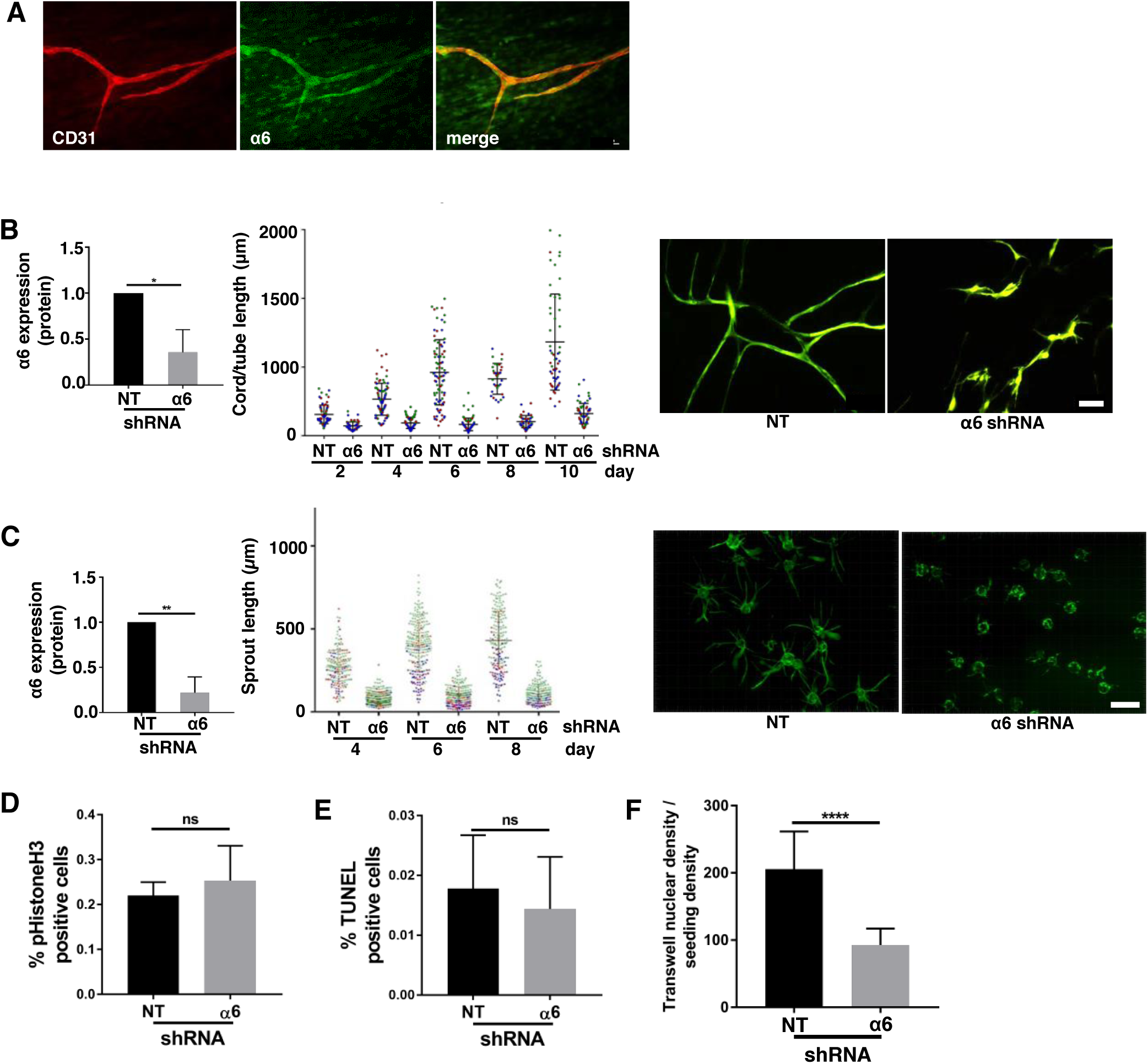
Depletion of endothelial α6 integrins inhibits endothelial tubulogenesis. **(A)** Immunofluorescence staining of 2-day planar co-cultures indicating the endothelial expression of α6 integrins. **(B)** Alpha6 integrins were depleted in endothelial cells by shRNA and the effects were assayed in 3 independent experiments. Efficiency of knockdown was assayed by western blot (left panel). The effect of α6 depletion on cord/tube length in planar co-culture was assayed every 2 days for 10 days and compared to control. Cord/tube length was quantified in three independent experiments. Plotted is the length of individual cords/tubes within each experiment represented by a different color (middle panel). Statistical analysis of knockdown was performed with a two-tailed Student’s t-test. The mean ± s.d is indicated. Representative images are shown from the 4-day co-cultures (right panel). Scale = 100 µm. **(C)** Similar effects of α6 knockdown were observed in bead sprout assays. Efficiency of α6 knockdown was determined by western blot (left panel). Sprout length was measured every 2 days for 8 days. Plotted is the length of individual sprouts from 3 independent experiments with each experiment represented by a different color (middle panel). Statistical analyses of knockdown was performed with a two-tailed Student’s t-test. The mean ± s.d. is indicated (left panel). Representative images of 6-day bead sprouts assays are shown (right panel). Scale = 500 µm. Proliferation **(D)** and survival **(E)** of cells expressing non-targeting (NT) or α6-targeting shRNA were analyzed by staining for phospho-histone H3 and TUNEL, respectively. Three fields were analyzed from each of three independent experiments. **(F)** Analysis of nuclear staining in transwell migration assays with non-targeting (NT) or α6-targeting cells. Five fields were analyzed from each of three independent experiments. Data was analyzed using two-tailed Student’s t-test. The mean ± s.d. is indicated. ns = no significance, *p ≤ 0.05, **p ≤ 0.01, ****p ≤ 0.0001.

The depletion of α6 integrins had a similar inhibitory effect in the bead sprout assay (Fig. 3C). Data was obtained from three independent experiments in which α6 expression was inhibited by the induction of α6 targeting shRNA (Fig. 3C, left panel). Measurements of sprout length overtime revealed that endothelial sprouting remained inhibited throughout the 8-day assay (Fig. 3C, middle panel). Images of representative bead sprouts formed at day 8 by control and α6 depleted endothelial cells are shown in Fig. 3C (right panel). The depletion of α6 integrins phenocopies the depletion of the LM-α5 chain and LM-α4 chain, suggesting that the interaction between α6 integrins and laminins secreted by endothelial cells is crucial for endothelial morphogenesis. Importantly, inhibition of these interactions, by depleting endothelial cells of α6 integrins did not significantly impact cell proliferation or survival (Fig. 3D&E), but did inhibit the migratory behavior of these cells on gelatin in a transwell migration assay (Fig 3F).

### Laminin-511 and α6 integrins regulate the expression of the pro-angiogenic genes CXCR4 and ANGPT2

To gain further insight into the mechanisms by which endothelial laminins and α6 integrins regulate tubulogenesis, we asked whether laminin-411, laminin-511 and α6 integrins regulate the same set of angiogenesis-associated genes. Since depleting endothelial cells of laminin-411 also inhibited sprouting, we tested whether expression of the same or a distinct set of genes was affected. We focused on the expression of genes previously associated with sprouting angiogenesis. These include VEGFR2, CXCR4, ANGPT2, Dll4, PDGFB, NRP1, JAG1, and MMP14 (De Smet et al., 2009; del Toro et al., 2010; Strasser et al., 2010). RNA was isolated from non-targeting and α6 targeting shRNA expressing endothelial cells and gene expression was analyzed by qPCR. Interestingly, depletion of either laminin-511 or α6 integrins led to significant decreases in RNA transcripts for CXCR4 and ANGPT2 (Fig. 4A&B). Additionally, a significant decrease in the expression of the LM-α5 chain RNA was also observed when α6 integrin expression was inhibited (Fig. 4A). Depleting endothelial cells of laminin-411 had little effect on the expression of these genes (Fig. 4C).

**Figure 4.**
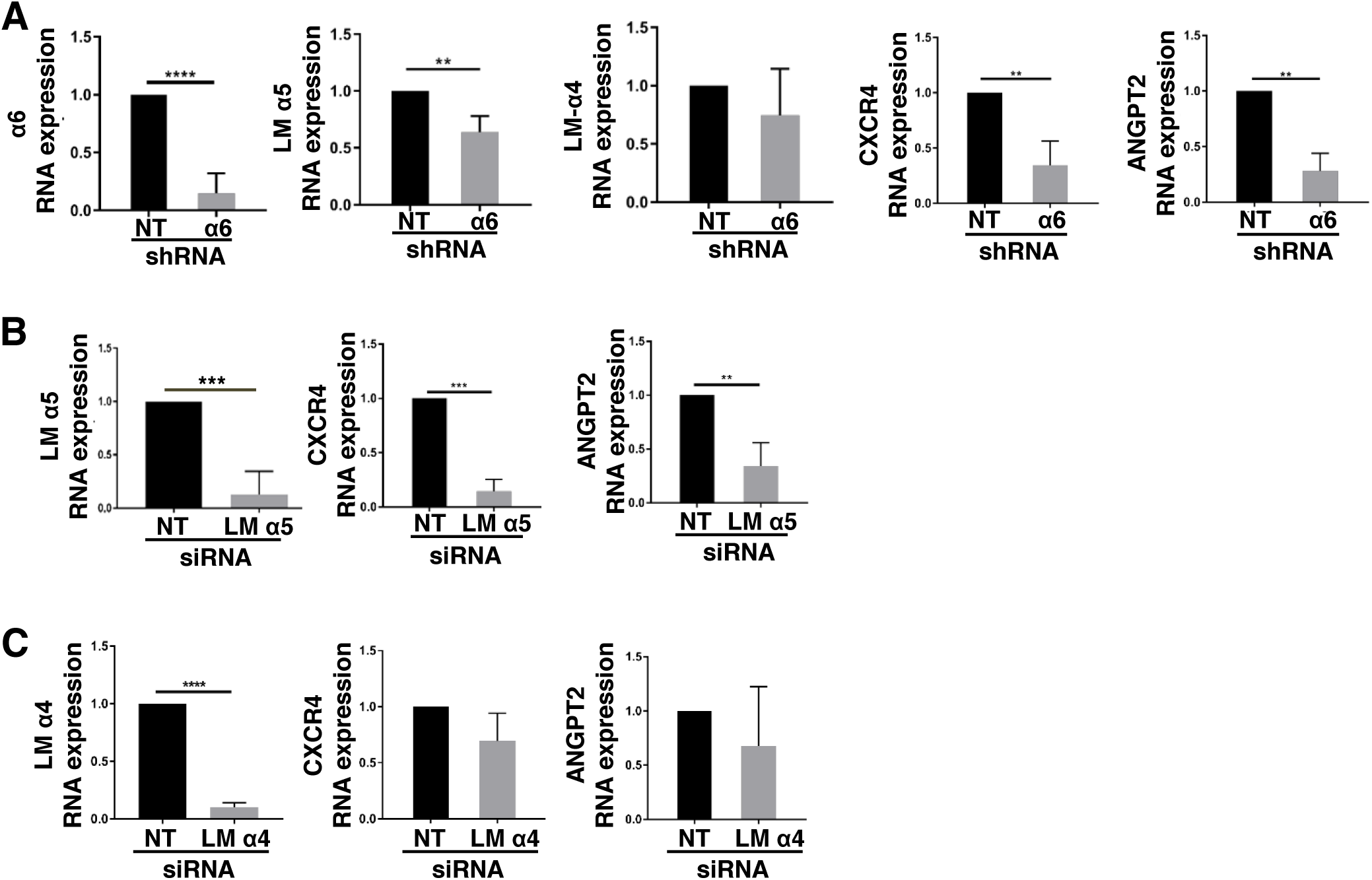
The expression of the CXCR4 and ANGPT2 genes are positively regulated by α6 integrins and laminin-511. **(A-C)** RNA was isolated from 6-day bead sprouts and analyzed by qPCR. **(A)** Shown is the efficiency of α6 depletion and the effects of α6 depletion on the expression of the LM-α4, LM-α5, CXCR4 and ANGPT2 normalized to non-targeting control. **(B)** Shown is the efficiency of depletion of the LM-α5 chain and the effects of this depletion on the expression of CXCR4 and ANGPT2. **(C)** Shown is the efficiently of depletion of the LM-α4 chain and the effects of this depletion of the expression of CXCR4 and ANGPT2. n = 3, **p ≤ 0.01, ***p ≤ 0.001, ****p ≤ 0.0001.

### CXCR4 signaling is required for endothelial morphogenesis

Since the best characterized mechanism of action for angiopoietin-2, the product of the ANGPT2 gene, is to antagonize the effects of angiopoietin-1 secreted by neighboring mural cells (Carmeliet and Jain, 2011), we did not analyze the role of ANGPT2 expression in our organotypic models. However, we did analyze the contribution of the chemokine receptor, CXCR4, as it has been previously implicated in vascular development and angiogenesis (Salcedo and Oppenheim, 2003; Salvucci et al., 2002; Tachibana et al., 1998; Unoki et al., 2010). Stromal-derived factor-1 (SDF-1) is a ligand for CXCR4, and is expressed by both endothelial cells (Salvucci et al., 2002) and fibroblasts (Nagasawa, 2014; Quan et al., 2015). Thus, it seemed possible that CXCR4 signaling is required downstream of α6 integrins and laminin-511 for endothelial morphogenesis in our models. To test this possibility, we used the pharmacological inhibitor AMD3100, which blocks CXCR4 activity (Hatse et al., 2002 65; Kalatskaya et al., 2009), as well as two different siRNA targeting sequences. In dose response experiments, AMD3100 inhibited both the number and length of sprouts (Fig. 5A). Significant effects on sprout numbers were observed at 500 nM, whereas sprout length was inhibited at concentrations as low as 100 nM (Fig. 5A). Interestingly, we did not need to add SDF-1 to the cultures as previously described (Strasser et al., 2010). Similarly, endothelial morphogenesis was inhibited in planar co-culture assays in a dose-dependent manner with significant inhibition starting at 100 nM of the inhibitor (Fig. 5B). Similar results were obtained when we inhibited the expression of CXCR4 using two different siRNAs. Both siRNA-targeting sequences significantly inhibited the expression of CXCR4 (Fig. 5C), as well as endothelial sprouting (Fig. 5D). As expected, depletion of either CXCR4 or α6 integrins inhibited CXCR4 protein expression (Fig. 5E). We next tested whether the expression of recombinant CXCR4 could rescue endothelial morphogenesis in α6-depleted cells. This was accomplished by transducing endothelial cells with a lentiviral vector expressing WT human CXCR4, which resulted in an increase of CXCR4 that was approximately an order of a magnitude higher than control (Fig. 5F). These cells were then depleted of α6 integrins and subsequently assayed for endothelial morphogenesis in three independent experiments. For each experiment, both the efficiency of α6 depletion and CXCR4 expression were measured by qPCR (Fig. 5G). The results indicated that the re-expression of CXCR4 partially rescued the α6-knockdown phenotype (Fig. 5H). Notably, the increased CXCR4 expression did not significantly impact endothelial morphogenesis in the control (Fig. 6H). These data indicate that the regulation of CXCR4 expression by α6 integrins and laminin-511 contributes to endothelial morphogenesis.

**Figure 5.**
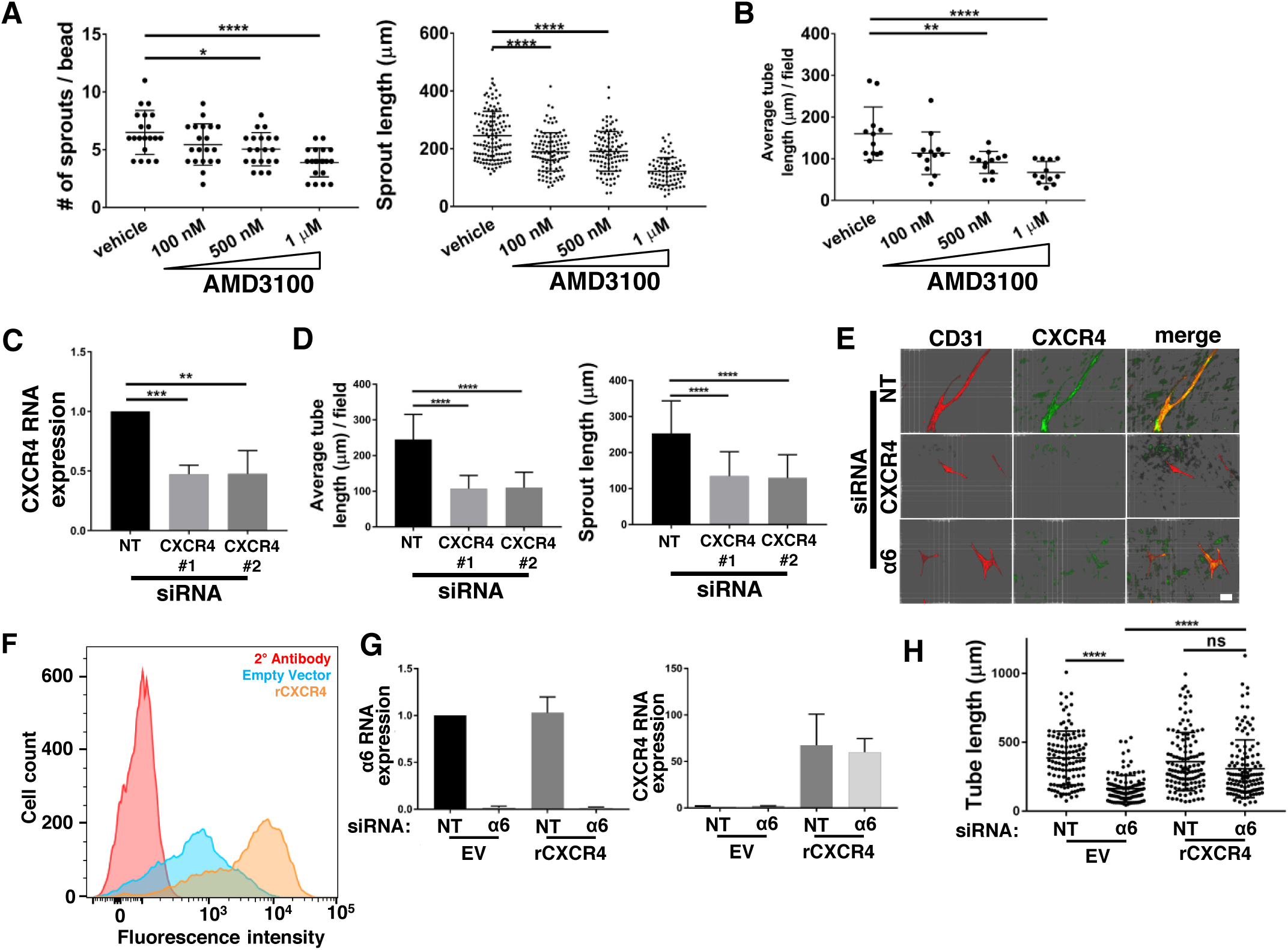
Endothelial CXCR4 is necessary for tubular morphogenesis in organotypic culture. **(A)** A dose-curve response using the specific CXCR4 inhibitor, AMD3100, shows a dose-dependent reduction in both number of endothelial sprouts, as well as sprout length in 6-day bead sprout assays. Left panel - Plotted is the number of sprouts on individual beads from 3 fields containing 7-10 beads from each of two independent experiments. Right panel-plotted is the length of individual sprouts on these same beads. Data was analyzed by one-way ANOVA and plotted as the mean length ± s.d. **(B)** AMD3100 inhibition of CXCR4 in 6-day planar co-culture causes defective morphogenesis in a dose-dependent manner. Six fields were analyzed in each of two independent experiments. Data was analyzed by one-way ANOVA and plotted as the mean length ± s.d. **(C)** The efficiency of CXCR4 depletion using 2 siRNA targeting sequences determined by qPCR. **(D)** Average length of tubes/field analyzed from 3 independent experiments with 6 randomly selected fields from each (Left panel). Individual sprout lengths were measured from 3 independent experiments with 10 randomly selected fields each, averaging 6-8 beads per field (Right panel). Data was analyzed by two-tailed Student’s t-test. **(E)** Immunofluorescence staining of 2-day planar co-cultures showing CXCR4 expression in non-targeting (NT), CXCR4-targeting and α6-targeting endothelial cells. Endothelial cells were stained with CD31 (red). Scale = 40 µm. **(F)** Expression of CXCR4 in endothelial cells transduced with empty vector (EV) or rCXCR4-expressing vector was measured by flow cytometry. **(G)** The efficiency of siRNA-mediated knockdown of α6 (Left Panel) and the RNA expression of CXCR4 (Right Panel) in CXCR4- and Empty Vector (EV)-expressing cells was assayed by qPCR for each of the three independent experiments. **(H)** Plotted are the lengths of individual tubes from 4-day planar co-cultures. Data was from 3 independent experiments with 5-6 randomly selected fields per experiment and analyzed by one-way ANOVA. ns = no significance, ****p ≤ 0.0001.

**Figure 6.**
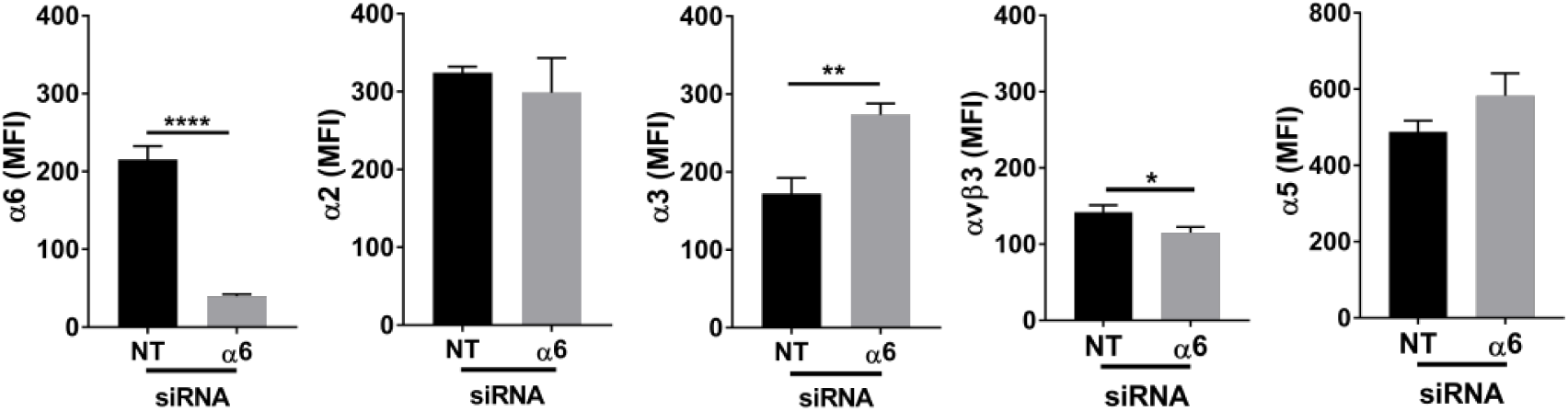
Depletion of α6 integrins alters the surface integrin profile. Flow cytometry analysis of the surface integrin expression of α2β1, α3β1, α5β1, and αvβ3 in endothelial cells transfected with non-targeting (NT) or α6-targeting siRNA. Plotted is the mean fluorescence intensity (MFI) ± s.d. from three independent experiments.

### Depletion of α6 integrins does not inhibit the expression of other integrins

To determine whether the depletion of α6 integrins affected the surface expression of other integrins known to bind to laminin-511 and/or have been implicated in angiogenesis (Avraamides et al., 2008; Davis and Senger, 2005; Halann et al., 2005; Kortesmaa et al., 2000; Song et al., 2017; Turner et al., 2017), we analyzed the surface expression of α2, α3, and α5 subunits and the αvβ3 integrin in cells treated with non-targeting or α6 targeting siRNA by flow cytometry and compared the mean fluorescence intensity from three independent experiments (Fig. 6). The data indicate that the surface expression of α2β1 and α5β1 was not significantly altered in cells depleted of α6 integrins. The surface expression of α3β1 was increased by almost 50%; however, this change did not compensate of the loss of α6 integrins in our functional assays (Fig. 3). The expression of αvβ3 was slightly but significantly decreased; however depletion of endothelial cells of αvβ3 did not inhibit endothelial morphogenesis on our organotypic model (Supplementary Figure 3).

### The expression of α6 integrins promotes the association of laminin-511 with endothelial tubes and promotes their integrity

Although we can detect laminin-511 at early stages of endothelial morphogenesis, to more easily determine whether depletion of α6 integrins inhibits the interaction of laminin-511 with endothelial cells, planar co-cultures were set up with endothelial cells that were transduced with lentiviruses carrying either α6-targeting or non-targeting shRNAs. Co-cultures were incubated in the absence of doxycycline for 10 days at which time normal endothelial tubes were well established. Co-cultures were then treated with doxycycline to induce the expression of non-targeting or α6-targeting shRNAs and imaged after 12 days of doxycycline treatment. Our results indicate that the expression of the laminin-511 is lost after the depletion of α6 integrins (Fig. 7A). In contrast, the laminin-411 α4 is still expressed, but appears diffuse compared to control (Fig. 7A). Interestingly, after the induction of α6 shRNA, endothelial tubes became increasingly destabilized over time exhibiting changes in cell shape and tube morphology (Fig. 7B). Endothelial cells became less elongated and tube morphology with in small patches endothelial monolayers, suggesting disruption of cell-cell junctions (Fig. 7C); however, no obvious changes in VE-cadherin expression were observed (Supplementary Fig 4). These results suggest that α6 integrins and laminin-511 promote endothelial tube integrity.

**Figure 7.**
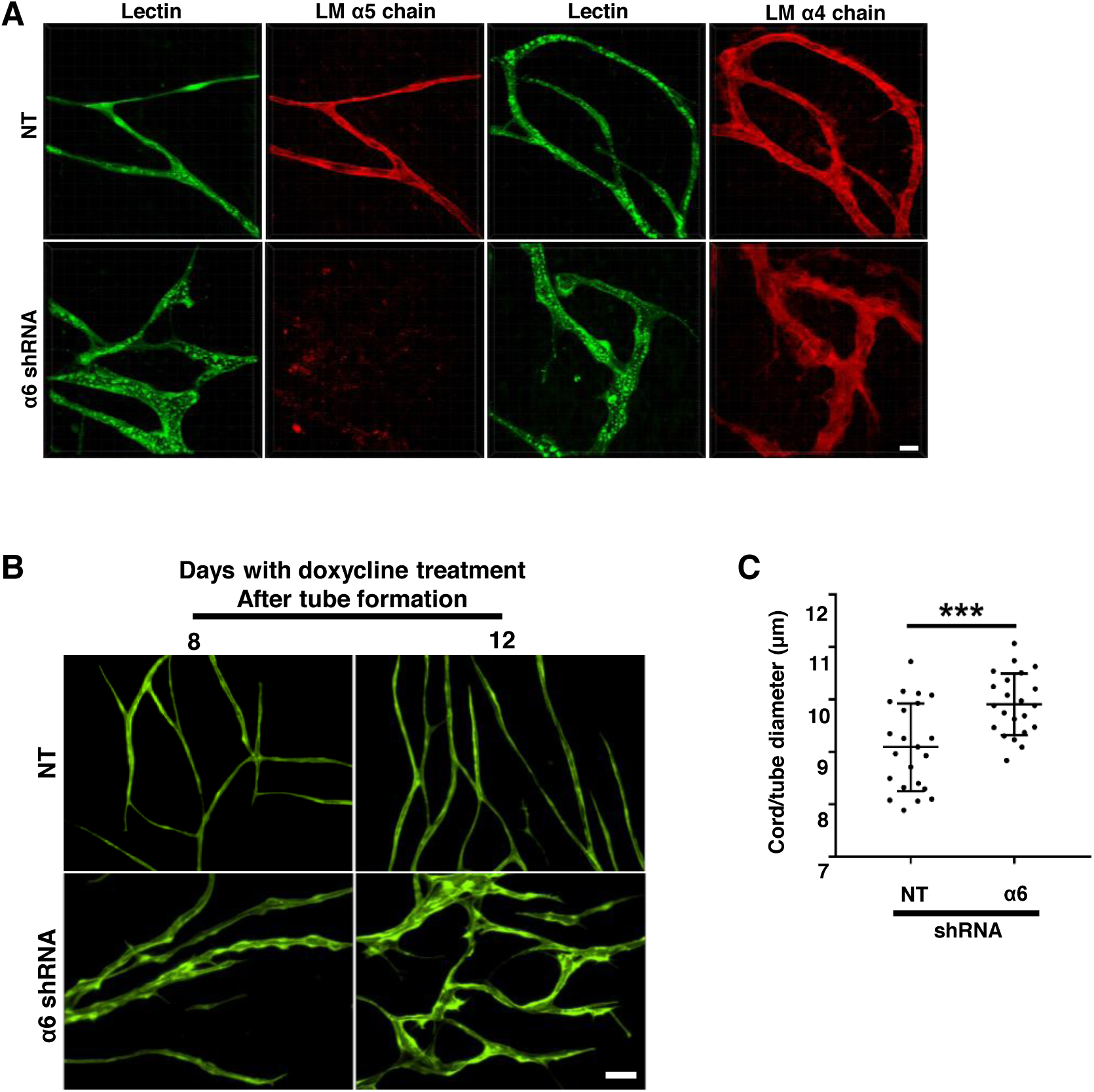
Depletion of α6 integrins from established endothelial tubes disrupts tube integrity and inhibits the expression of laminin-511. **(A-C)** Endothelial tubes were formed for 10 days in the planar co-culture assay and the expression of α6 or non-targeting (NT) shRNA was induced with doxycycline for 12 days. **(A)** Expression of laminin-511 and laminin-411 in endothelial tubes after the depletion of α6 integrins compared to control. Scale = 25 µm. **(B)** Representative images of endothelial tubes after 8 and 12 days of induction of either the α6 or non-targeting (NT) shRNA. Scale = 100 µm. **(C)** Plotted is the average tube width per field in (NT-sh) and (α6-sh) structures measured from ten randomly selected fields in two independent experiments. n=22. Student’s t-test was used to compare conditions. The mean ± s.d. is indicated. ***p ≤ 0.001.

## Discussion

Using organotypic co-culture angiogenesis assays, we demonstrated that endothelial secretion of laminin-411 and laminin-511, and the expression of α6 integrins that bind to these laminins are required for endothelial morphogenesis. In these assays, laminin-411 and laminin-511 did not compensate from one another. As an approach to understand the underlying mechanisms, we analyzed the expression of a number of genes that are regulated during angiogenesis, some of which are enriched in tip cells of sprouting vessels (De Smet et al., 2009; del Toro et al., 2010; Strasser et al., 2010). The majority of genes interrogated did not change upon depletion of either the LM-α4 chain, the LM-α5 chain or the integrin α6 subunit. Interestingly, CXCR4, and ANGPT2 expression were inhibited in cells depleted of either LM-α5 chain or the integrin α6 subunit, but not in cells depleted of the LM-α4 chain, suggesting that α6 integrins and laminin-511 promote angiogenesis by regulating the expression of these genes. We further investigated the contribution of CXCR4 and employing a number of experimental approaches demonstrated that CXCR4 functions downstream of α6 integrins during endothelial tubulogenesis.

We detect changes in CXCR4 and ANGPT2 RNA, suggesting that α6 integrins and laminin-511 impacted either the transcription of these genes or the stability of their RNA transcripts. Integrins are known to regulate gene expression by these mechanisms (For examples, see Iyer et al., 2005; Streuli and Akhtar, 2009) and one or both mechanisms may be involved in regulating the expression of CXCR4 and ANPT2. Multiple transcription factors have consensus motifs in the promoters for these two genes. The presence of consensus motifs for the TEAD family of DNA binding proteins is intriguing and suggests that the YAP/TAZ transcriptional co-activators are potential candidates for the regulation of CXCR4 and ANGPT2 during endothelial morphogenesis. Promoters for Dll4 and KDR, which are not regulated by α6 integrins and laminin-511, do not contain consensus motifs for TEAD binding. Importantly, laminins and their integrin receptors are known to activate YAP/TAZ in a number of physiological contexts (Chang et al., 2015; De Rosa et al., 2019; Elbediwy et al., 2016; Hu et al., 2017; Zhang et al., 2017).

There are two isoforms of the integrin α6 subunit, the more common α6A isoform and the α6B isoform (de Melker and Sonnenberg, 1999). In human breast cancer stem cells, the α6Bβ1-dependent adhesion to laminin-511 activates the transcriptional co-activator, TAZ, which in a feed forward loop promotes the expression of the α5 chain of laminin-511 (Chang et al., 2015). It is unclear whether endothelial cells express the α6B isoform. Nonetheless, we found that depletion of α6 integrins from endothelial cells results in a significant, but relatively small decrease in LM-α5 RNA. This decrease is unlikely responsible for the dramatic decrease we observe in laminin-511. It is possible that α6 integrins regulate the translation or processing the LM-α5 chain or the secretion of laminin-511. It is notable that we do not detect significant levels of the LM-α5 chain intracellularly by immunofluorescence microscopy.

In contrast to the depletion of laminin-511, there was no overlap between the genes regulated by loss of expression of the laminin-411 and the loss of α6 integrins. We anticipated that the expression of Notch ligand, DLL4 would be inhibited upon LM-α4 depletion, as previous studies demonstrated that DLL4 expression is down regulated in LM-α4 null mice during developmental retinal angiogenesis (Stenzel et al., 2011). It is possible that other integrins, in addition to α6 integrins, engage laminin-411 to promote different aspects of endothelial morphogenesis. The α2β1 integrin is a potential contributor as previous *in vitro* studies demonstrated that α2β1 plays a central role in endothelial cell adhesion to laminin-411 (Stenzel et al., 2011). Consistent with the down regulation of DLL4, LM-α4-null animals also exhibited hyper sprouting in the retina (Stenzel et al., 2011). Increased podosome formation was also observed in aortic ring angiogenesis assays (Seano et al., 2014). The Tie-2 dependent deletion of α6 integrins inhibited the formation of podosomes in these same assays, implicating these α6 integrins in this process (Seano et al., 2014). However, we did not observe hyper sprouting with the depletion of LM-α4 in our co-culture models. It is possible that the complex *in vivo* environment minimizes the requirement for laminin-411 in endothelial morphogenesis or that our organotypic co-culture models are more dependent upon endothelial laminin-411 expression.

Previous studies identified roles for laminin-411 and laminin-511 in promoting vessel stability (Thyboll et al., 2002; Song et al., 2017). Our results indicate that the loss of the expression of α6 integrins from established tubes results in the loss of laminin-511, but not laminin-411, and that the loss of LM-511 and α6 integrins correlates with the loss of tube integrity. VE-cadherin is known to regulate endothelial barrier function (Garrett et al., 2017), and previous studies indicate that LM-511 and β1 integrins promotes the recruitment of VE-cadherin to cell-cell junctions to promote vessel stability (Song et al., 2017; Yamamoto et al., 2015). In preliminary studies, we did not detect obvious changes in the expression of VE-cadherin in endothelial cells depleted of α6 integrins; however, it seems likely possible that the loss of α6 integrins and laminin-511 perturbs cell-cell adhesion.

The α6 subunit can dimerize with either the β1 or β4 subunit to form either the α6β1 or α6β4 integrin. Unlike α6β1, the expression of α6β4 is restricted to a subset of endothelial vessels (Desai et al., 2013; Hiran et al., 2003). We demonstrated that α6β4 is not expressed in angiogenic vessels during cutaneous wound repair or in sprouting vessels in explant angiogenesis assays, but becomes expressed during vessel maturation, after which its expression is restricted to a subset of vessels (Desai et al., 2013; Hiran et al., 2003). Other studies suggest that α6β4 is critical in signaling the onset of angiogenesis, but that its expression is down regulated in the newly forming vessels (Nikolopoulos et al., 2004). In brain endothelium, the α6β4 integrin was found to promote arteriolar remodeling in response to hypoxia and vascular integrity in response to autoimmune inflammation (Welser et al., 2017; Welser-Alves et al., 2013). It will be interesting in future studies to determine whether α6β4 contributes to endothelial morphogenesis and tube integrity in the organotypic co-culture models and if so to identify the mechanisms involved.

In summary, our data support the conclusion that α6 integrins and laminin-511 positively regulate the expression of CXCR4 to endothelial tubulogenesis and that laminin-411 and laminin-511 play distinct roles in this process.

## Materials and methods

### Cell Culture

Human umbilical vein endothelial cells (HUVECs) were from Lonza (Allendale, NJ) and were cultured in in EGM-2 (Lonza, CC-3162). Adult human dermal fibroblasts (HDFs) were isolated and characterized as previously described (Varney et al., 2016; Zheng et al., 2019) and generously provided by the Van De Water laboratory (Albany Medical College). Human embryonic kidney epithelial 293FT cells (HEK293FT) were a kind gift from Dr. Alejandro Pablo Adam lab (AMC). HDFs and HEK293FT cells were cultured in DMEM (Sigma D6429) containing 10% FBS (Atlanta Biologicals), 100 units/ml penicillin (Life Technologies), 100 µg/ml streptomycin (Life Technologies), and 2.92 µg/ml L-glutamine (GE LifeSciences). All cells were cultured at 37°C in 5% CO2.

### Antibodies and Reagents

Antibodies used in this study were from Dako (Santa Clara, CA), Cell Signaling (Danvers, MA), Santa Cruz Biotechnology Inc. (Santa Cruz, CA), Sigma Aldrich (St. Louis, MO), BD Biosciences (Billerica, MA), Abcam (Cambridge, MA), MilliporeSigma (Burlington, MA), and ThermoFisher Scientific (Waltham, MA). Additional antibody information can be found in Table 1.

**Table 1:** Antibodies

| **Protein** | **Host** | **Manufacturer** |
| --- | --- | --- |
| CD31 [JC70A] | Mouse | Dako |
| CD31 [M-20] | Rabbit | Santa Cruz |
| FITC-conjugated UEA lectin (L9006) |  | Sigma Aldrich |
| Collagen IV (ab6586) | Rabbit | Abcam |
| Laminin-111 (ab11575) | Rabbit | Abcam |
| Laminin α4 chain [3H2] (ab205568) | Mouse | Abcam |
| Laminin α5 chain [4B12] (ab77175) | Mouse | Abcam |
| CXCR4 (ab1670) | Goat | Abcam |
| CXCR4 [EPUMBR3] (ab181020) | Rabbit | Abcam |
| Phospho-Histone H3 (Ser10) | Rabbit | Cell Signaling |
| α6 integrin subunit [GoH3] | Rat | BD Biosciences |
| α6 integrin subunit [F-6] | Mouse | Santa Cruz |
| α2 integrin subunit [P1E6] | Mouse | Millipore |
| α3 integrin subunit [P1B5] | Mouse | Millipore |
| α5 integrin subunit [P1D6] | Mouse | Millipore |
| αvβ3 integrin [LM609] | Mouse | Millipore |
| VE-Cadherin (D87F2) | Rabbit | Cell Signaling |
| GAPDH (GA1R) | Mouse | ThermoFisher Scientific |
| Anti-rat IgG Alexa Fluor 488 (A21208) | Donkey | ThermoFisher Scientific |
| Anti-mouse IgG Alexa Fluor 488 (A21202) | Donkey | ThermoFisher Scientific |
| Anti-mouse IgG Alexa Fluor 568 (A10037) | Donkey | ThermoFisher Scientific |
| Anti-rabbit IgG Alexa Fluor 488 (A11034) | Goat | ThermoFisher Scientific |
| Anti-mouse HRP (ab97046) | Rabbit | Abcam |

### siRNA

HUVECs were plated in 6-well tissue culture plates and transfected with siRNA at a 50 nM concentration with RNAiMAX (ThermoFisher) using the protocol provider by the manufacturer. HUVECs transfected with siRNA were assayed for knockdown and used in planar co-cultures (described below) 48 h after transfection. HUVECs transfected with siRNA and used in bead sprout assays (described below) were transfected during bead coating and assayed for knockdown at the end of experiment. Additional siRNA information can be found in Table 2.

**Table 2:**
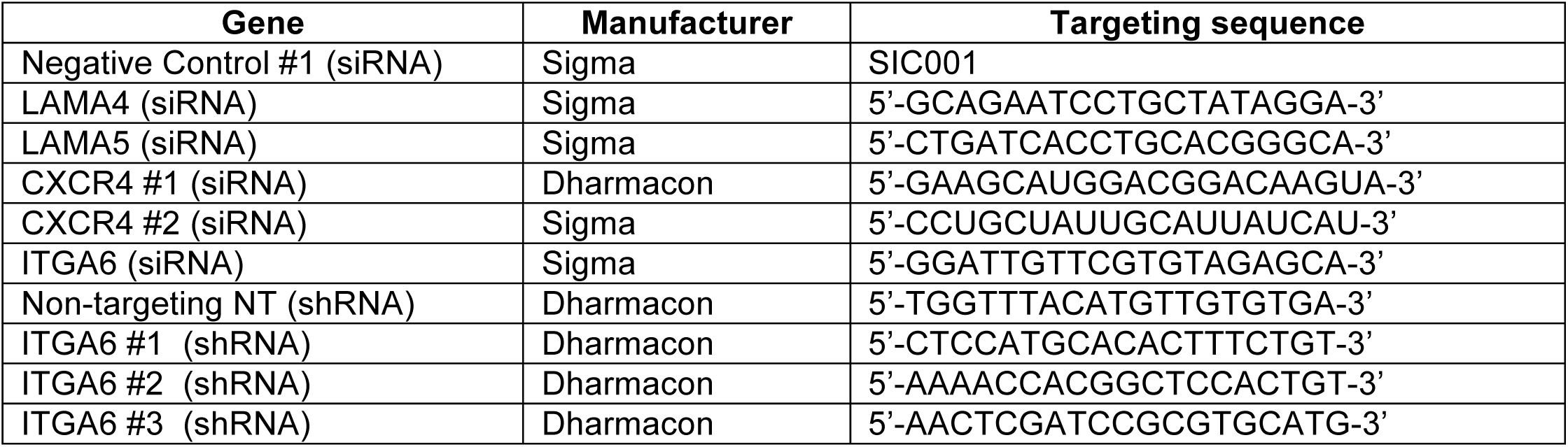
RNAi

### Inducible shRNA

Doxycycline-inducible lentiviral (SMART) vectors harboring shRNAs targeting the α3, α6, β4 integrin subunit or a non-targeting (NT) shRNA were purchased from Dharmacon (Lafayette, CO). Lentiviruses were produced by co-transfection of HEK293FT cells with the shRNA expression vector together with the packaging plasmid, psPAX2, coding for Gag, Pol, Rev, Tat (#12260, Addgene), and the envelope plasmid, pMD2.G, coding for VSV-G (#12259, Addgene). HUVECs were transduced with filtered viral supernatant plus 8 µg/ml polybrene. Cells were induced with doxycycline (100 ng/ml) for 48 h prior to adding HUVECs to co-cultures or in some experiments after 10 days in co-culture as described below. Additional shRNA information can be found in Table 2.

### Exogenous CXCR4 expression

Human WT CXCR4 was acquired as a donor plasmid (#81957, Addgene) and recombined into a lentiviral destination vector (#25890, Addgene). Lentiviruses were produced by co-transfection of HEK293FT cells with the CXCR4 expression vector together with the packaging plasmid, psPAX2, coding for Gag, Pol, Rev, Tat (#12260, Addgene), and the envelope plasmid, pMD2.G, coding for VSV-G (#12259, Addgene). HUVECs were transduced with filtered viral supernatant plus 8 µg/ml polybrene.

### Quantitative PCR (qPCR)

TRIzol (ThermoFisher) was used to isolate RNA from siRNA transfected HUVECs, as well as, shRNA expressing HUVECs. Extraction of RNA from bead sprout assays (described below) using TRIzol was performed after the removal of HDFs with trypsin-EDTA Solution 10X (59418C, Sigma). cDNA was synthesized with iScript Reverse Transcription Supermix (BioRad) using 1 µg of RNA. Equal amounts of cDNA were used in qPCR reactions performed with iQ SYBR Green Supermix (BioRad). Values were normalized to the signal for β-actin. The nucleotide sequences of the qPCR primers used are listed in Table 3.

**Table 3:**
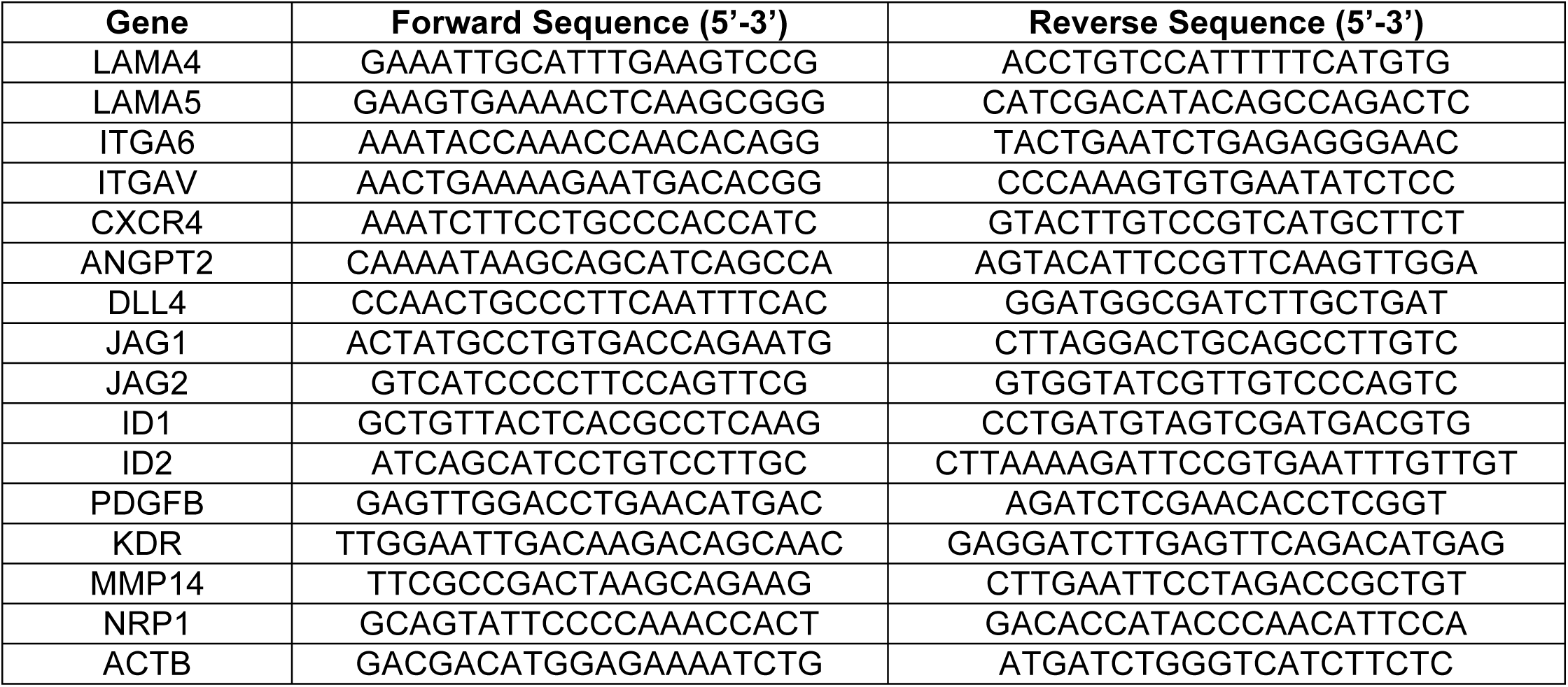
qPCR primers

### Flow cytometry

Analysis of cell surface integrin expression was performed by staining live cells with primary antibodies (1:100 dilution) for 1 hour on ice. Cells were washed with ice-cold PBS 2X and stained with secondary antibodies (1:1000 dilution) for 30 minutes on ice. Cells were fixed with 2% PFA after 2X washes with ice-cold PBS. Analysis of rCXCR4 expression was performed by fixing cells with 4% PFA in PBS for 5 minutes and permeablizing with 0.5% Triton X-100 for an additional 5 minutes. Cells were washed with PBS 2X and treated with Calf Intestinal Alkaline Phosphatase (CIP) for 1 h at 37°C prior to staining with primary and secondary antibodies. Data was acquired with a FACSCalibur (BD Biosciences) and analyzed using FlowJo.

### Migration

SMART Vector shRNA expressing cells were induced with doxycycline (100 ng/ml) for 48 h prior to assays. Cells were cultured in serum-free EGM-2 medium overnight prior to assay. Fifty thousand cells were seeded in triplicates into transwells and a separate 24-well plate. Serum-containing EGM-2 was then added to the lower chamber of transwells followed by incubation for 4 hours at 37°C in 5% CO2. Transwells were fixed with 4% PFA (Electron Microscopy Sciences) and stained with DAPI. The lower membrane was imaged with a 4X objective and density quantified using ImageJ (NIH). Cell seeding efficiency was determined by performing toluidine blue assays in the 24-well plates. These assays were performed by fixing cells with 70% ethanol at room temperature for 1 hour, followed by wash with dH2O and staining with 0.05% toluidine blue at room temperature for an additional 2 hours. After wash with dH2O, toluidine blue was extracted with 10% acetic acid at 0.3 ml / well and absorbance measured at 650 nm, using 405 nm as reference on a Synergy2 microplate reader (BioTek Instruments). An empty well was processed the same way and used for baseline. Migration efficiency was determined by dividing DAPI density by absorbance.

### Proliferation and survival

SMART Vector shRNA expressing cells were induced with doxycycline (100 ng/ml) for 48 h and re-seeded for overnight culture. Cells were fixed with 4% PFA (Electron Microscopy Sciences) for 5 minutes and permeabilized with 0.5% Triton X-100 in PBS for an additional 5 minutes. After 2X wash with PBS, a TUNEL assay was performed using DeadEnd Fluorometric TUNEL System (Promega) following manufacturer’s protocol. Cells were then blocked with 2% BSA and stained for Phospho-Histone H3 (1:100 dilution) (Cell Signaling) in 2% BSA overnight at 4°C. After 3X washes for 1 hour, cells were incubated with DAPI and secondary antibodies (1:1000 dilution) for 1 hour at room temperature. Images were acquired with a Plan Fluor 10X/0.30 objective and analyzed using ImageJ (NIH).

### Western blotting

Western blotting was used to confirm RNAi induced knockdown. Cells were lysed in mRIPA buffer (50 mM Tris pH 7.4, 1% NP-40, 0.25% Na Deoxycholate, 150 mM NaCl, 1 mM EDTA) containing both phosphatase (Sigma, #4906837001) and protease inhibitor cocktails (ThermoFisher, 78440). Equal amounts of protein (20 to 40 µg) were separated by SDS-PAGE and transferred to nitrocellulose for antibody probing. Imaging was performed with a ChemiDoc XRS+ (BioRad) and quantitation with Image Lab (BioRad).

### Organotypic culture assays

#### Planar co-culture

As one organotypic culture, we utilized the planar co-culture model developed by Bishop and colleagues (Bishop et al., 1999) and modified by the Pumiglia lab (Bajaj et al., 2012). This model reconstitutes some of the complex interactions that occur during angiogenesis among endothelial cells, the ECM and supporting cells. To set up the co-culture, HDFs were seeded in tissue cultures dishes ± glass coverslips and cultured to confluence. The medium was changed to EGM-2. HUVECs, expressing targeting or non-targeting RNAi, were then seeded 16 h later at a density of 20,000 cells per 9 cm^2^ and cultured up to 10 days. ShRNA-mediated knockdown in pre-formed tubes was accomplished by culturing HUVECs expressing doxycycline-inducible lentiviral (SMART) vectors on HDFs for 10 days, in the absence of doxycycline with medium changed every 48 h. Doxycycline was then introduced on day 11, at a concentration of 100 ng/ml and refreshed every 48 h for up to 16 days. Endothelial morphogenesis and changes in tube structure were analyzed by immunofluorescence microscopy.

#### Bead Sprout assay

To study endothelial sprouting, we employed the bead sprout assay as described by Nakatsu and Hughes (Nakatsu and Hughes, 2008). Cytodex 3 beads (GE) were coated at ∼1000 HUVECs per bead inside of a 2 ml microcentrifuge tube for 4 at 37°C, mixing gently by inverting the tubes every 20 min and transferred to a T25 flask and incubated at 37°C, overnight. Beads were then washed 3X with EGM-2 medium and re-suspended in PBS containing 3 mg/ml of fibrinogen (Sigma, #F8630) and 0.15 U/ml of aprotinin (Sigma, #A6279). Thrombin (Sigma, #T4648) was added at a final concentration of 0.125 U/ml and the mixture was plated in wells of and 8-well slide (Corning, #3-35411). The mixture was allowed to clot for 30 min at 37°C. HDFs were then added to the top surface of the fibrin gel in EGM-2 medium at a concentration of 30,000 cells per well. The formation of sprouts and sprout lengths were assayed by either immunofluorescence of phase contrast microscopy.

### Immunofluorescence Microscopy

Planar co-cultures were fixed with 4% PFA (Electron Microscopy Sciences) for 15 min, permeabilized with 0.5% Triton X-100 in PBS for 15 min, and then blocked with 2% BSA in PBST for 1 h at RT. Antibodies were diluted in 2% BSA in PBST and incubated with cells overnight at 4°C. Samples were then washed 4X with PBST at RT over the course of 4 h, and then incubated for 1 h with the appropriate secondary antibodies (1:1000 dilution). Following secondary antibody staining, samples were washed 3X with PBST at RT for 1 h and mounted with SlowFade Gold antifade reagent (ThermoFisher). Samples were analyzed using a Nikon inverted TE2000-E microscope equipped with phase contrast and epifluorescence, a digital CoolSNAP HQ camera, a Prior ProScanII motorized stage and a Nikon C1 confocal system and EZC1 and NIS-Elements acquisition software. Images were acquired with Plan Fluor 4X/0.13, Plan Fluor 10X/0.30, Plan Fluor ELWD 20X/0.45, Plan Apo 40X/1.0 oil, and Plan Apo 100X/1.4 oil objectives and analyzed with either NIS elements (Nikon) or IMARIS software as indicated. Contrast and/or brightness were adjusted for some images to assist in visualization. Fibroblasts were removed from bead sprout assays using trypsin-EDTA Solution 10X (59418C, Sigma) prior to fixation and staining. Samples were analyzed using a Zeiss LSM 880 inverted confocal microscope running under Zeiss ZEN2.3 software and equipped with AiryScan detector, as well as 2 standard photomultiplier detectors in combination with a 32 channel ultra-sensitive GaAsP detector. Images were acquired with 10X/0.45 NA and Plan-Apochromat 63X/1.4 NA oil DIC objectives and analyzed with IMARIS software.

### Image analysis

For planar co-cultures, where endothelial cells initially form cell trains/cord, which mature into endothelial tubes, cord/tube lengths in planar co-cultures were measured by tracing tubes within each field using NIS Elements (Nikon). Any tubes that extend beyond the field were excluded from analysis. Tube widths in planar co-cultures were calculated with AngioTool (NIH) by dividing total tube area by total tube lengths per field. Average tube length per field was calculated with AngioTool (NIH). Sprout lengths in bead sprout assays were measured by tracing each sprout using NIS elements (Nikon) and sprouts per bead were counted manually.

### Statistical analysis

Statistical analysis was performed with GraphPad Prism software using Student’s t-test or one-way ANOVA. P value of p<0.05 was considered to be statistically significant.

## Acknowledgements

The authors thank Drs. Livingston Van De Water and C Michael DiPersio for critically reading this manuscript, Dr. Van De Water for providing human adult dermal fibroblasts, Debbie Moran for assistance in the preparation of the figures and the Imaging Core of Albany Medical College.

## Data availability

All reagents generated in this study will be made available upon request.

---

**Supplemental Figure 1.**
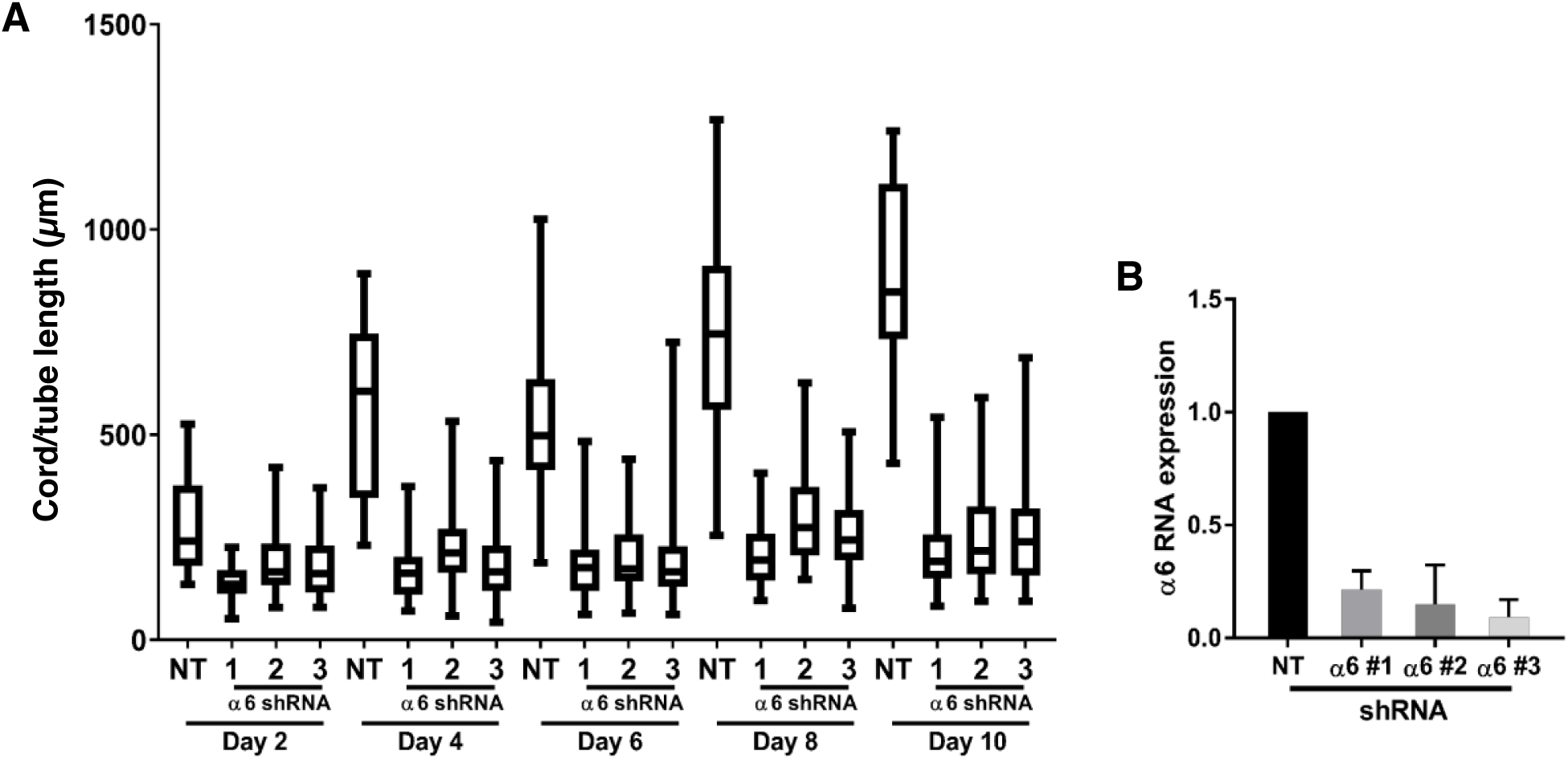
**(A)** Depletion of endothelial α6 integrins using 3 independent shRNA targeting sequences shows similar inhibition of tubular morphogenesis compared to control. Cord/tube lengths were collected from 10 randomly selected fields in 3 independent experiments. **(B)** Knockdown of α6 was confirmed using qPCR. Mean RNA expression is from 3 independent experiments. Alpha 6 targeting sequence 2 was used in Figures 3, 4, and 7 of main text.

**Supplemental Figure 2.**
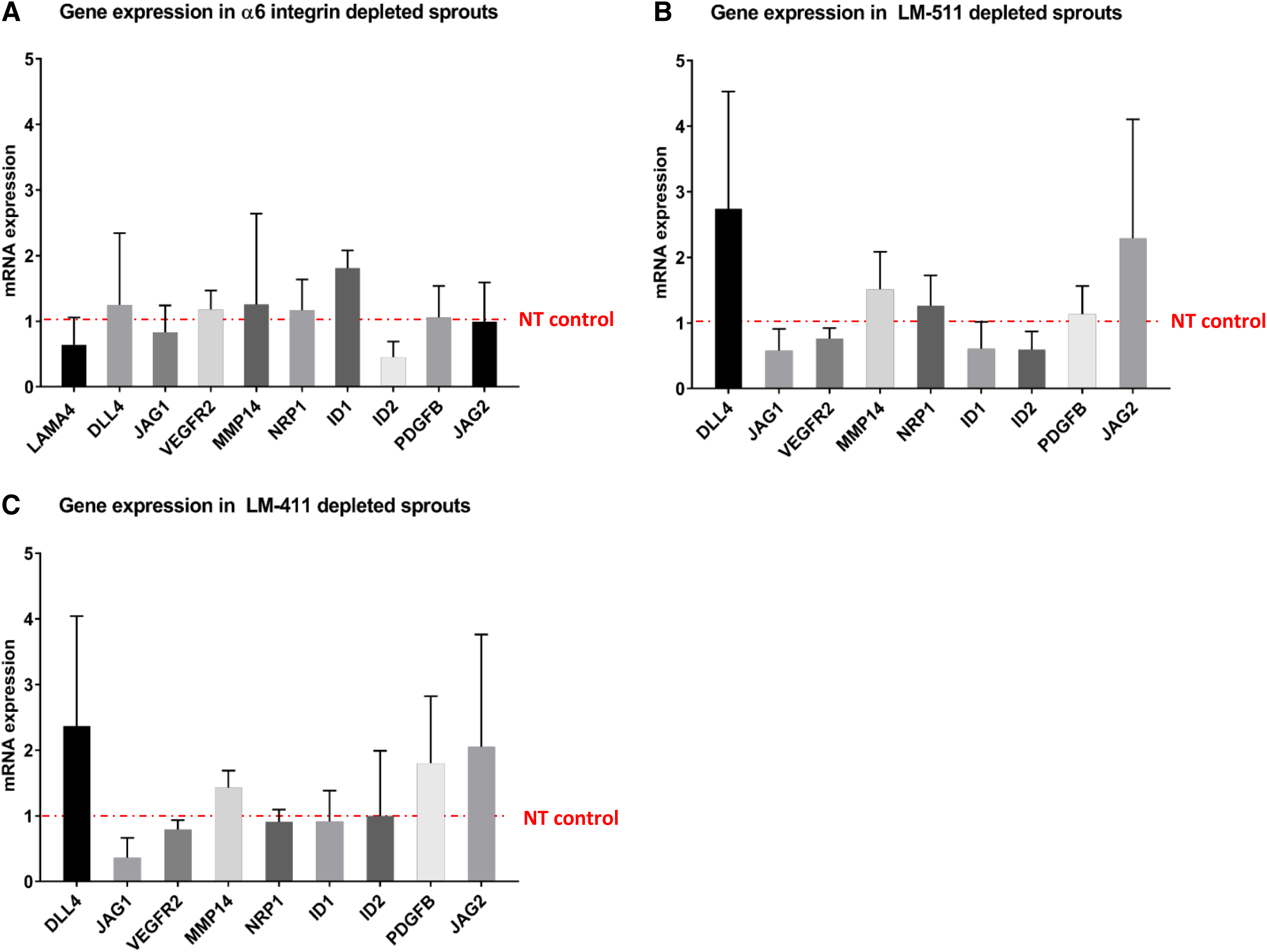
Expression of angiogenesis-associated genes in 5-day bead sprout assay compared to control **(A)** alpha 6 integrin-depleted endothelial cells **(B)** laminin-511-depleted endothelial cells and **(C)** laminin-411-depleted endothelial cells were measured by qPCR. Data are plotted as the mean ± s.d. from 3 independent experiments.

**Supplemental Figure 3.**
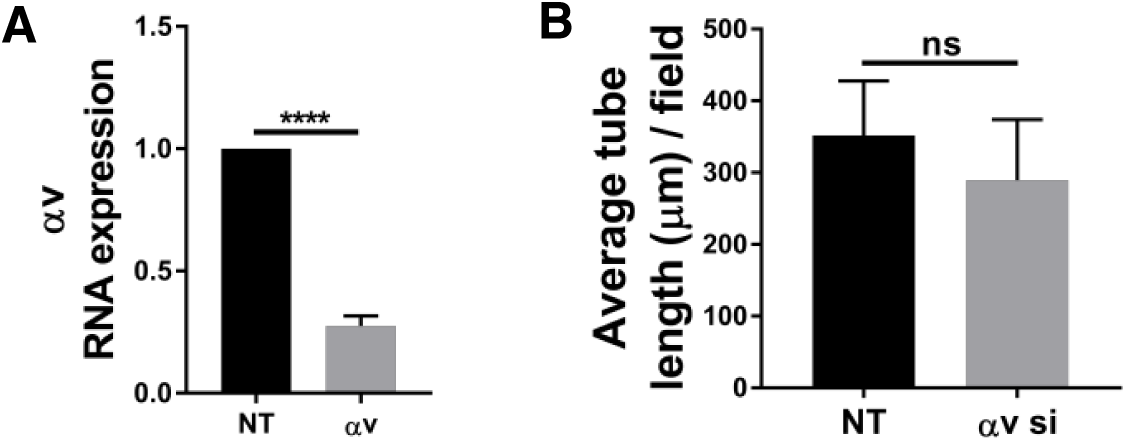
**(A)** Efficiency of αv depletion quantified by qPCR and **(B)** average tube length at day 4 compared to control in planar co-culture.

**Supplemental Figure 4.**
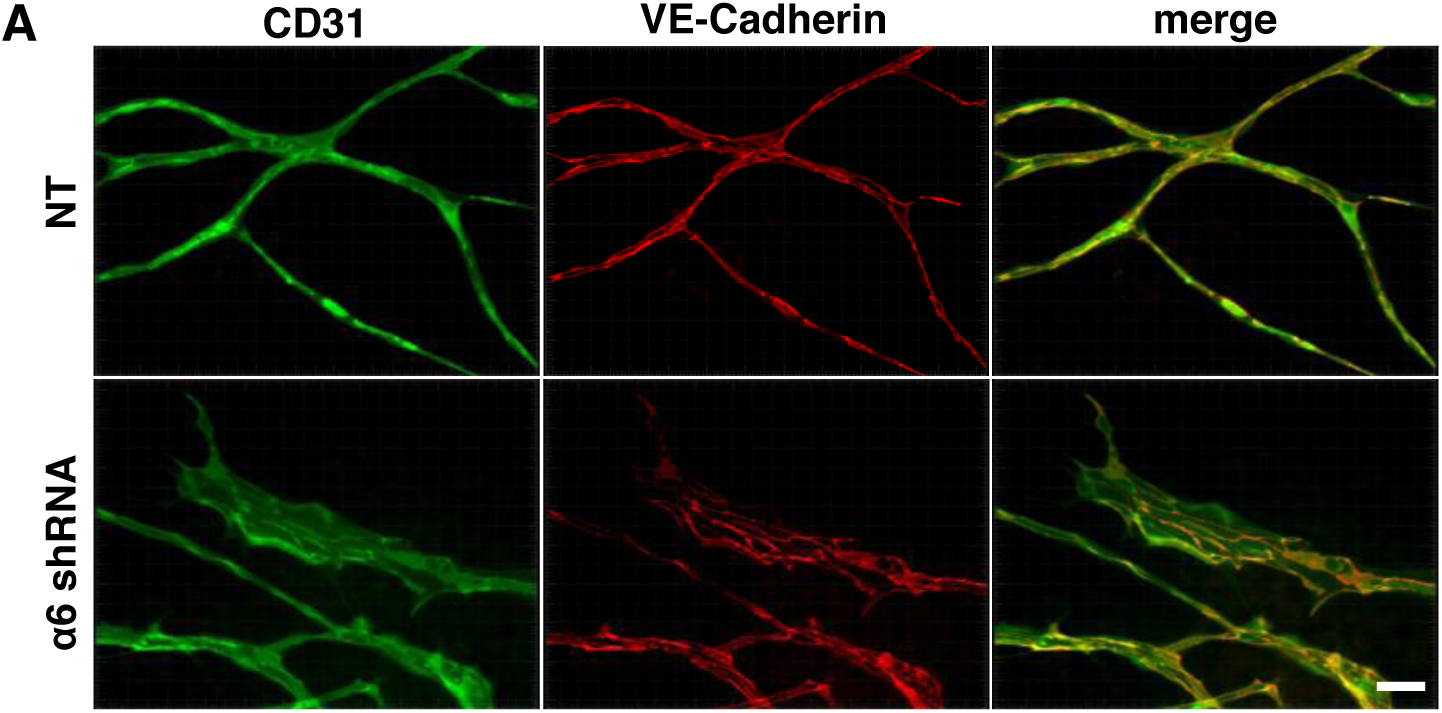
**(A)** Endothelial structures after 12 days of doxycycline induction in non-targeting (NT) or α6-targeting shRNA stained for CD31 and VE-Cadherin. Scale = 50 µm.

